# High Frequency Oscillations (250-500Hz) in Animal Models of Alzheimer’s Disease and Two Animal Models of Epilepsy

**DOI:** 10.1101/2022.06.30.498284

**Authors:** Christos Panagiotis Lisgaras, Helen E. Scharfman

## Abstract

**Objective:** To test the hypothesis that high frequency oscillations (HFOs) between 250 and 500Hz occur in mouse models of Alzheimer’s disease (AD) and thus are not unique to epilepsy.

**Methods:** Experiments were conducted in three mouse models of AD: Tg2576 mice that simulate a form of familial AD, presenilin 2 knock-out (PS2KO) mice, and the Ts65Dn model of Down’s syndrome. We recorded HFOs using wideband (0.1-500Hz, 2kHz) intra-hippocampal and cortical surface EEG at 1month until 24months-old during wakefulness, slow wave sleep (SWS) and rapid eye movement (REM) sleep. Interictal spikes (IIS) and seizures were also analyzed for the possible presence of HFOs. Comparisons were made to the intra-hippocampal kainic acid and pilocarpine models of epilepsy.

**Results:** We describe for the first time that hippocampal and cortical HFOs are a new EEG abnormality in AD mouse models. HFOs occurred in all transgenic mice but no controls. They were also detectable as early as 1month of age and prior to amyloid-β plaque neuropathology. HFOs were most frequent during SWS (vs. REM or wakefulness). Notably, HFOs in the AD and epilepsy models were indistinguishable in both spectral frequency and duration. HFOs also occurred during IIS and seizures in the AD models, although with altered spectral properties compared to isolated HFOs.

**Significance:** Our data demonstrate that HFOs, an epilepsy biomarker with high translational value, are not unique to epilepsy and thus not disease specific. Our findings also strengthen the idea of hyperexcitability in AD and its significant overlap with epilepsy. HFOs in AD mouse models may serve as an EEG biomarker which is detectable from the scalp and thus amenable to non-invasive detection in people at risk for AD.

**KEY POINTS:**

- High frequency oscillations (HFOs, 250-500Hz) occur in mouse models of Alzheimer’s disease
- HFOs are detectable from the hippocampus and overlying cortex
- HFOs are most frequent during slow wave sleep
- HFOs in AD mouse models resemble HFOs in two animal models of epilepsy
- HFOs can be detected during interictal spikes and seizures in the AD models

## INTRODUCTION

Alzheimer’s disease (AD) affects a growing number of individuals worldwide, placing a tremendous burden on their lives and the lives of their caretakers.^1^ Regrettably, there are few treatments and many fail to resolve symptoms, indicating the need for identifying additional targets for therapeutic intervention.^2^

One of the contributors to AD pathophysiology is abnormal electrical activity,^3^ which is often underestimated since the use of EEG in AD is limited. Notably, seizures in patients with AD^4^ are known to accelerate disease progression^5^ and neuronal cell loss.^6^ Independently of seizures, patients with AD also show interictal spikes (IIS) which negatively affect cognition.^7^ In this context, anti-seizure medications (ASMs) can improve cognitive function in those AD patients with IIS.^8^ ASMs were also effective in amnestic patients before a late stage of AD, suggesting that abnormal electrical activity is involved early during disease progression.^9^ Therefore, a better understanding of abnormal electrical activity in AD is potentially important from a mechanistic and therapeutic perspective.^10^

In epilepsy, abnormal electrical activity includes IIS and seizures as well as HFOs. HFOs are implicated in seizure onset^11^ and used as a biomarker of the seizure onset zone.^12^ Motivated by epilepsy studies, we expanded the recording bandwidth and sampling rate of conventional EEG^13^ and asked whether HFOs, defined by a spectral frequency between 250 and 500Hz, can be detected in the hippocampus and cortex of mouse models of AD. To that end, we recorded from 3 mouse models of AD, each with a different causal mechanism. In addition, we implemented the intra-hippocampal kainic acid (IHKA) and pilocarpine (PILO) models of epilepsy to determine possible commonalities of HFOs in the AD mouse models and HFOs occurring in epilepsy.^14,15^ EEG recordings were made from the dentate gyrus (DG) because of past work showing it contributes to cognitive and electrical abnormalities in AD mouse models.^16^ EEG recordings were analyzed for the presence of HFOs during all behavioral states (wakefulness, SWS, REM) using methods previously validated in human^17^ and rodent^15^ EEGs.

HFOs with a spectral frequency of 250 to 500Hz were recorded from the DG of all transgenic mice and not in the controls. Notably, HFOs were also detectable from the frontal cortical contact. The number of HFOs in all transgenic mice was significantly elevated during SWS relative to other behavioral states, consistent with abnormal SWS in humans with AD.^18^ HFOs in the AD mouse models were indistinguishable from HFOs in animal models of epilepsy both in spectral frequency and duration. We also identified HFOs during IIS and seizures in the AD mouse models, and these HFOs were distinct from those occurring at other times. Our data provide new insight into AD pathophysiology and suggest that HFOs could serve as a new EEG biomarker for AD. Because cortical surface recordings showed HFOs, wideband-EEG could become a non-invasive screening tool in people at risk for AD.

## MATERIALS AND METHODS

### I. Animals

Experiments were conducted in 3 mouse models of AD. Mice carrying a mutation (Tg2576+/-, PS2KO+/-, Ts65Dn) are termed transgenic and wild type littermates (Tg2576-/-, PS2KO+/+, 2n) are termed controls. Tg2576 mice (n=5 controls; n=8 transgenic) overexpress the human amyloid precursor protein (APP) using the hamster prion promoter and have the Swedish mutation (APP_Swe_).^19^ Tg2576 transgenic mice develop memory deficits^20^ and amyloid-β deposits by 6months and robust plaque pathology by 12months of age.^19,21^ Also, Tg2576 mice show IIS and seizures,^20^ which allowed us to determine if HFOs coincide with them. Tg2576 transgenic mice were 5.5±2.4months old (range 1-19months) at the time of recording, when IIS and seizures are intermittent.^20^ Tg2576 control mice were 5.3±3.1months-old (range 1-18). No statistically significant age differences were found between control and transgenic Tg2576 mice (Mann-Whitney U-test, U=18, p=0.80). Two additional Tg2576 transgenic mice were used for seizure monitoring and their age is shown in Supplemental table 1. Presenilin 2 knock-out (PS2KO) mice (n=4 controls; n=4 transgenic) lack the catalytic subunit of γ-secretase, critical to APP metabolism to amyloid-β.^22^ PS2KO mice were used because mutations in the *PS2* gene are an AD precipitant in familial early-onset AD cases.^23^ In addition, PS2KO transgenic mice do not develop amyloid-β neuropathology^24^ which allowed us to study HFOs independent of it. PS2KO mice were 10.7±1.9months old (range 6-15months) at the time of recording. PS2KO control mice were 10.7±1.9months-old (range 6-15months). No statistically significant age differences were found between control and transgenic PS2KO mice (unpaired Student’s t-test, t=0.009, df=6, p=0.99).

The Ts65Dn model (n=5 controls; n=4 transgenic) of Down’s syndrome have a portion of the mouse chromosome 16 containing the *APP* gene triplicated.^25^ Ts65Dn mice show profound basal forebrain cholinergic neuronal degeneration late in the disease which also occurs in human AD.^25^ Ts65Dn transgenic mice were 15.6±3months old (range 3-24months) at the time of recording.

Ts65Dn control mice were 15.3±4.6months-old (range 3-24months). No statistically significant differences were found between control and transgenic Ts65Dn mice (Mann-Whitney U-test, U=20.5, p=0.82). In addition, there were no statistically significant differences in age between the controls of different mouse lines (Kruskal-Wallis test, H(3)=3.89, p=0.14) and no differences in transgenics of the different mouse lines (Kruskal-Wallis test, H(3)=4.84, p=0.08). The age of all the mice used in the study are shown in Supplemental table 1.

To compare HFOs in the AD mouse models with HFOs recorded in animal models of epilepsy (n=6 IHKA; n=4 PILO) we used 8wk old C57BL/6J controls of *Amigo2*-Cre+/- mice (#027, Charles River Laboratories). IHKA- and PILO-treated mice were recorded at 3-4months after IHKA or PILO treatment.

All mice were bred on a C57BL/6J background and fed Purina 5001 (W.F. Fisher), with water *ad libitum*. Cages were filled with corn cob bedding and there was a 12hr light:dark cycle (7:00a.m. lights on, 7:00p.m. lights off). Genotyping was performed by the Mouse Genotyping Core Laboratory at NYU Langone Medical Center or in-house. All experimental procedures were performed in accordance with the NIH guidelines and approved by the Institutional Animal Care and Use Committee at the Nathan Kline Institute.

### II. Epilepsy models

We used the IHKA model as described previously.^15^ In brief, mice were anesthetized with 3% isoflurane (Henry Schein) and then transferred to a stereotaxic apparatus (World Precision Instruments). One burr hole was drilled above the left hippocampus (—2mm anterior-posterior (A-P) to Bregma, -1.25mm medio-lateral (M-L)). A 0.5ml Hamilton syringe (#7001, Hamilton) was lowered from the brain surface 1.6mm into the left hippocampus (—1.6mm D-V) and 100nL of 20mM KA was injected over 5min. The needle remained in place for an additional 5min to prevent backflow and the incision was closed with Vetbond (3M). Next, mice were monitored for *status eilpeticus* (SE) defined as severe and continuous convulsive^26^ seizures lasting for several hrs.^15^ Mice were implanted with electrodes 4wks after IHKA, when they already show seizures.^15^

A separate cohort of mice was treated with PILO. Mice were injected with the muscarinic antagonist scopolamine methyl nitrate (1mg/kg s.c., Sigma Aldrich) to reduce the peripheral effects of PILO. The β2-adrenergic agonist terbutaline hemisulfate (1mg/kg s.c., Sigma Aldrich) was injected to support respiration. Ethosuximide (150mg/kg s.c., Sigma Aldrich) was administered to reduce the occurrence of brainstem seizures which often lead to mortality. SE was induced by injecting pilocarpine hydrochloride (250mg/kg, s.c., Sigma Aldrich) and after 2hrs, mice were injected with diazepam (5mg/kg, s.c., Hospira) to reduce SE severity and 1mL (s.c.) of lactated Ringer’s (Aspen Veterinary Resources) for hydration. PILO-treated mice were implanted with electrodes 4wks after PILO.

### III. Electrode implantation

Mice were anesthetized with 3% isoflurane and then transferred to a stereotaxic apparatus where anesthesia was maintained with 1-2% isoflurane. Next, buprenorphine (0.05mg/kg, s.c.) was injected to reduce discomfort. Two burr holes were drilled above the cerebellar region and subdural screw electrodes (2.54mm length stainless steel) were placed and stabilized using dental cement (Lang Dental) to serve as reference (—5.7mm A-P, +1.25mm M-L) and ground (—5.7mm A-P, -1.25mm M-L).

One burr hole was drilled over the left hippocampus (—1.9mm A-P, -1.2mm M-L). For electrodes, we used either a single wire electrode (90μm California Fine Wire) or a 16channel silicon probe (#A1×16, Neuronexus or #PLX-QP-3-16E-v2, Plexon) to ensure that HFOs would not be missed. Electrodes were placed in the left dorsal DG (—1.9mm A-P, -1.2mm M-L, -1.9mm D-V). The position of the electrode was confirmed post-mortem with Nissl stain. For bilateral DG recordings, a second electrode was implanted into the right DG (—1.9mm A-P, +1.2mm M-L, -1.9mm D-V). For simultaneous recordings in the DG and cortex, a single wire was implanted in the DG and a screw electrode over the right frontal cortex (—0.5mm A-P, +1.5mm M-L).

### IV. Wideband electrophysiological recordings

For recording, mice were housed into a 21cm × 19cm square transparent plexiglass cage with access to food and water. A pre-amplifier (Pinnacle Technologies or 16-channel headstage with accelerometer, Intan) was connected to the implant and then to a rotatory joint (Pinnacle Technologies or Doric Lenses) via a lightweight cable to allow unrestricted movement of the mouse.

EEG signals were amplified 10x and recorded at 2kHz sampling rate using a bandpass filter (0.1-500Hz) with Sirenia Acquisition software. These recording settings are used to capture HFOs in the 250-500Hz frequency range.^13^ For mice implanted with a silicon probe we used either the Digital Lynx SX (Neuralynx) or an RHD interface board (Intan) using the same settings as above. This allowed 16channel signals to be sampled simultaneously. Although the 3 recording systems had different gains, when raw data were normalized, the amplitude of the recorded signals were not significantly different across the 3 recording systems (Sirenia: 1.7±0.10μV, n=15; Intan: 1.6±0.34μV, n=9; Neuralynx: 2.6±0.33μV, n=3; Kruskal-Wallis test, H(3)=4.17, p=0.12). High frame rate video (>30frames/sec) was recorded simultaneously using an infrared camera (ac2000, Basler or CCTV, Apex). Mice were continuously recorded for 3 consecutive days at a minimum. For seizure monitoring, video-EEG recorded continuously for 1wk.

### V. HFO, IIS and seizure detection

HFOs were defined as oscillations >250Hz as our prior study of epileptic mice^15^ and consistent with human HFOs.^14,27,28^ We analyzed HFOs during wakefulness, SWS and REM sleep as before.^15^ An artifact-free 10min period was used to detect if HFOs occurred, consistent with clinical HFO identification.^13,29^ When HFOs were detected in a given mouse, they were present from one 10min period to the next, and in control mice without HFOs, all 10min periods showed 0 HFOs (Supplemental figure 5). These data suggested that 10min is a sufficiently long period and HFOs are stable enough so that one 10min period can be used to examine HFOs (see also^13,29^). For each mouse, three 10min periods (one during wakefulness, one during SWS, and one during REM) were sampled and HFOs/min were calculated for each mouse.

For detecting HFOs, we used an automated approach in the RippleLab application written in MATLAB^30^ and the algorithm developed by Staba and colleagues.^17^ In brief, each channel was band-pass filtered between 250-500Hz using a finite impulse response digital filter and the root mean square (RMS) amplitude of the filtered signal was quantified. Successive RMS amplitudes >5x standard deviations (SDs) above the mean amplitude of the baseline RMS signal, and a duration >6ms^31^ were collected for manual review. Baseline signals were calculated in the same behavioral state and same animal. Thus, each animal had its own baseline for detecting HFOs in each behavioral state that HFOs were examined (wakefulness, SWS, REM). We accepted a “putative” HFO as a “real” HFO based on established inclusion/exclusion criteria.^15,17,31^ Putative HFOs that had at least 4 peaks in the rectified band-passed signal >3xSDs greater than the mean of the baseline signal were included. Putative HFOs associated with artifactual events, movement or other sources of noise were excluded.^31^ Artifacts were identified by the absence of well-resolved central frequency in the time-frequency domain. Thus, artifacts showed an elongated “blob” appearance in the spectrogram; robust spectral power occurred throughout the time-frequency domain, as previously shown.^32^ Artifacts were excluded from the analysis.

To better appreciate HFO power in time, we applied time-frequency analyses to visualize HFO power. To that end, we used the time-frequency function which is part of RippleLab^30^ and applied a 64-point window to visualize HFO power in the 250-500Hz frequency.

For IIS detection we followed the same criteria as previously described.^20^ In brief, we included IIS that exceed REM’s background activity by >5xSDs above the mean during REM. Only REM sleep was analyzed for the presence of IIS because based on our prior study,^20^ IIS in Tg2576 mice are most robust during REM sleep and variably occur in other behavioral states besides REM. Seizures in the Tg2576 mice were detected using seizure detection criteria described in our prior study.^15^

### VI. Histology

Mice were deeply anesthetized with isoflurane and then injected with an overdose of urethane (2.5g/kg, i.p., Sigma Aldrich). Mice were transcardially perfused with 10ml of room temperature saline, followed by 30ml of cold 4% paraformaldehyde (Sigma Aldrich). The brain was quickly removed and stored overnight in 4% paraformaldehyde. Coronal brain sections (50μm) were cut using a vibratome (Ted Pella) and stored in cryoprotectant solution at 4°C until use.

For Nissl stain, sections were mounted on 3% gelatin-coated slides and allowed to dry overnight. Then slides were dehydrated in increasing concentrations of ethanol (70%, 95%, 100%, 100%) for 2.5min each, cleared in Xylene (Sigma Aldrich), and dehydrated again (100%, 100%, 95%, 70%) followed by hydration in double-distilled (dd) H_2_0 for 30sec. Then sections were stained with 0.25% cresyl violet (Sigma Aldrich) in ddH_2_0 for 1.5min followed by 30sec in 4% acetic acid. Next, sections were dehydrated (70%, 95%, 100%, 100%), cleared in Xylene, and cover-slipped with Permount (Electron Microscopy Systems). Electrode placement was imaged using a microscope (#BX51, Olympus of America) and digital camera (Infinity3-6URC) at 2752×2192 pixel resolution and Infinity capture software for micrographs.

### VII. Statistics

Data are presented as a mean ± standard error of the mean (SEM). Dots indicate individual data points. Data in the text are stated as a mean ± SEM. Statistical significance was set at p<0.05 and is denoted by asterisks on all graphs. Statistical comparisons that did not reach significance are not designated by a symbol in graphs but are reported in text.

All statistical analyses were performed using Prism (Graphpad). To determine if data fit a normal distribution, the Shapiro-Wilk test was used. Welch’s test addressed homogeneity of variance. Comparisons of parametric data of two groups were conducted using unpaired or paired two-tailed Student’s t-tests. When data did not fit a normal distribution, non-parametric statistics were used. The non-parametric test to compare two groups was the Mann-Whitney U-test. For comparisons of >2 groups, one-way ANOVA was used when data were parametric and Kruskal-Wallis for non-parametric data. When a statistically significant main effect was found by ANOVA, Bonferroni *post-hoc* tests were used with corrections for multiple comparisons and for Kruskal-Wallis, Dunn’s *post-hoc* tests were used. For parametric data with multiple comparisons, two-way ANOVA was used. When a significant effect of a main factor was found, Tukey’s *post-hoc* tests were used, with corrections for multiple comparisons. For correlation analyses we used linear regression. The Pearson correlation coefficient (r) was used for parametric data and Spearman r for non-parametric data.

## RESULTS

### I. HFOs are robust in the dentate gyrus of 3 AD mouse models and absent in controls

#### Tg2576 mice

Tg2576 mice were implanted with an electrode targeting the DG and electrode placement was verified post-mortem using Nissl-stained sections (Fig 1A and Supplemental figure 2). We first recorded from control mice and found no HFOs in any of the behavioral states (Fig 1B; example is from SWS). In contrast, Tg2576 transgenic mice showed robust HFOs (Fig 1C). Fig 1D shows HFOs during the trough of slow waves in the raw and filtered EEG traces. When the total number of HFOs was compared, after pooling data from all behavioral states, Tg2576 mice showed a significantly greater number of HFOs compared to controls (Mann-Whitney U-test, U=0, p=0.001; Fig 1E). Recordings with silicon probes suggested that HFOs are well localized to the DG and not recorded from overlying area CA1. A representative recording of HFOs in the sublayers of the DG and overlying area CA1 is shown in Supplemental figure 6. The results were similar for 7 silicon probe recordings from 5 transgenic mice (n=2 Tg2576, n=2 PS2KO, n=1 Ts65Dn).

**Figure 1:**
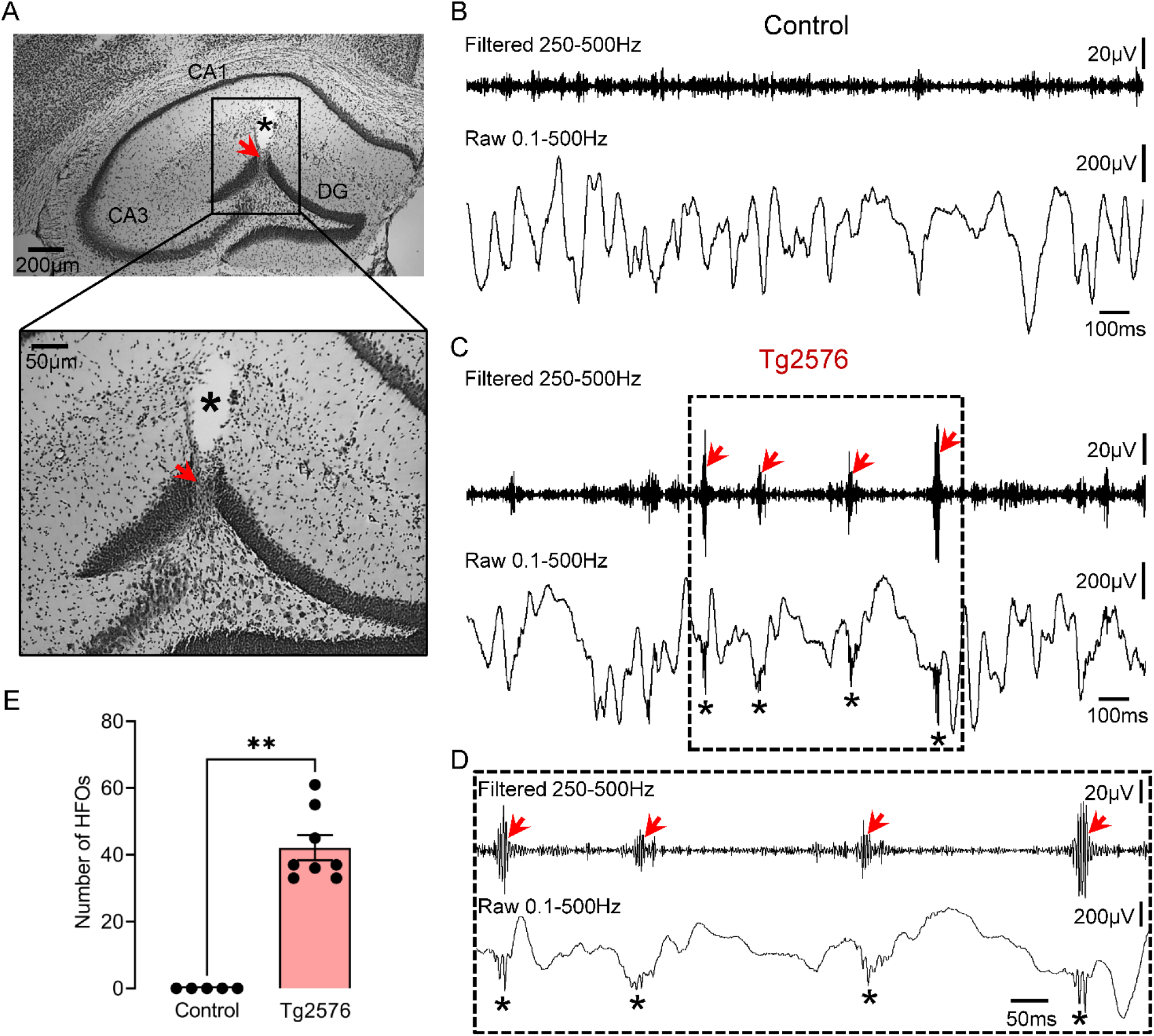
HFOs in the dentate gyrus of Tg2576 mice. (A) A representative Nissl-stained section confirming electrode placement in the dentate gyrus (DG; red arrow). The asterisk marks a cavity after electrode removal. Electrode targeting for both the control and transgenic mice is provided in Supplemental figure 2. (B) A representative EEG recording from the DG of a control mouse during SWS. Top trace is bandpass-filtered for 250-500Hz. Bottom trace is the raw (unfiltered) trace. (C) Same as in B but for a Tg2576 transgenic mouse. Note the presence of HFOs in the filtered trace (red arrows) and absence of HFOs in controls. HFOs were also visible in the raw trace at the troughs of slow wave activity (asterisks). (D) Expanded traces from the inset in C are shown. Note that HFOs (red arrows) stand out from baseline in the filtered trace. (E) Total number of HFOs (all behavioral states pooled) in control and Tg2576 transgenic mice. The total number is based on three 10min recordings one from each behavioral state (wakefulness, SWS, REM), and units are /min. HFOs were significantly more frequent (Mann-Whitney U-test, U=0, p=0.001) in Tg2576 transgenic mice (n=8) compared to control mice (n=5), where they were absent.

**Figure 2:**
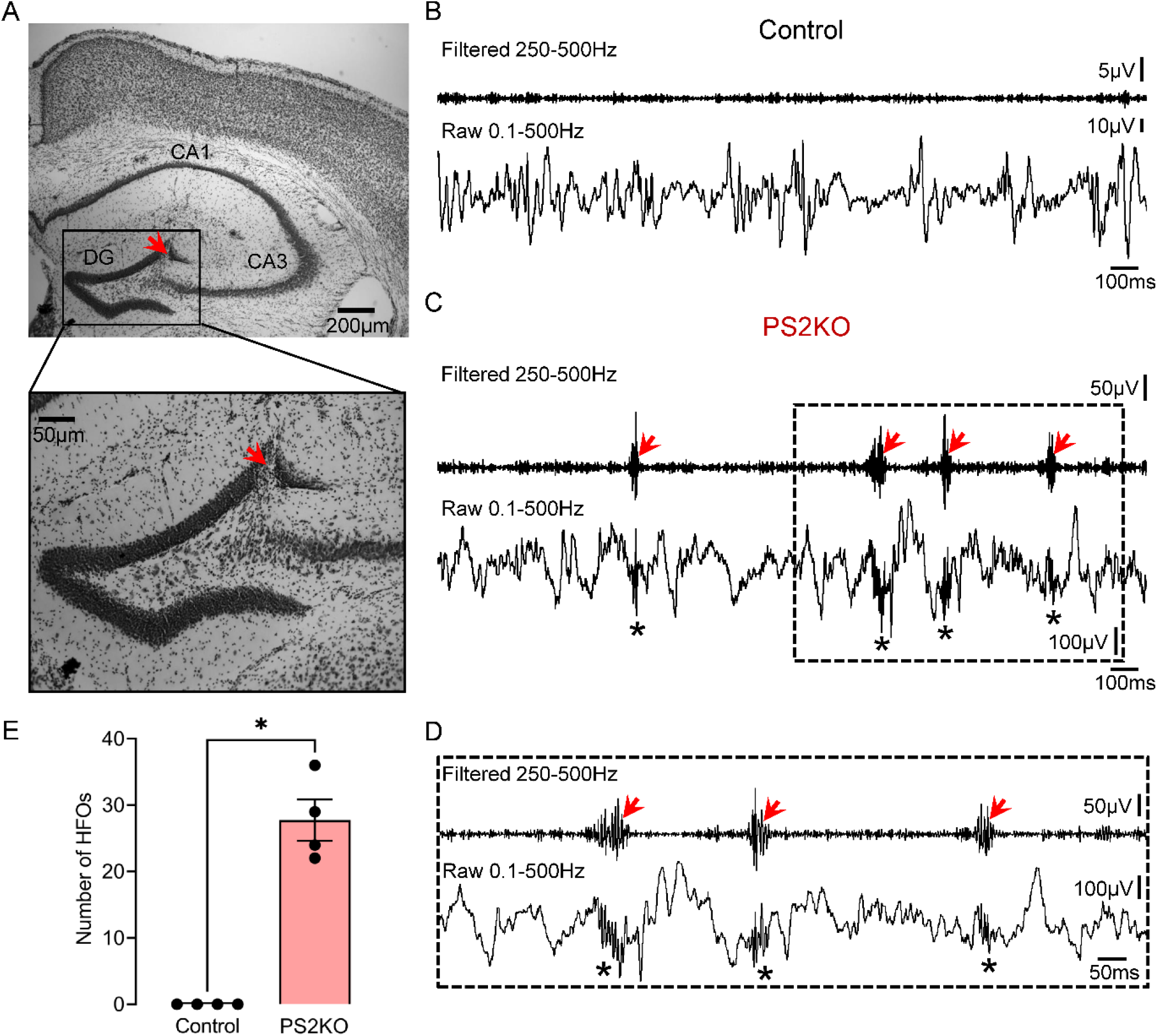
HFOs in the dentate gyrus of PS2KO mice. (A) A representative Nissl-stained section confirming electrode placement in the DG (red arrow). Electrode targeting for both the control and transgenic mice is provided in Supplemental figure 3. (B) A representative EEG recording from the DG of a control mouse during SWS. Note the absence of HFOs in filtered and raw traces. (C) Same as in B but the mouse was a PS2KO transgenic. Note the presence of HFOs in the filtered (red arrows) and raw traces (asterisks). Please note that the difference in amplitude between PS2KO transgenic and control is because mice were recorded using different recording systems. As we explain in Methods, comparisons were possible across recording systems since normalized data were not significantly different across systems. Please see Methods for more details. (D) Expanded traces from the inset in C are shown. Note that HFOs (red arrows) stand out from baseline in the filtered trace. (E) Total number of HFOs recorded from control (n=4) and PS2KO transgenic (n=4) mice. Data from all behavioral states were pooled. The total number of HFOs was significantly higher in PS2KO vs. control mice (Mann-Whitney U-test, U=0, p=0.02).

#### PS2KO and Ts65Dn mice

To determine generalizability of the Tg2576 mouse model, we recorded from 2 additional models. PS2KO transgenic mice had robust HFOs (Fig 2C to D). No HFOs were recorded in the 4 PS2KO control mice (Fig 2B). PS2KO transgenic mice showed a significantly greater number of HFOs compared to controls (Mann-Whitney U-test, U=0, p=0.02; Fig 2E). In the third AD mouse model, the Ts65Dn mouse, we also recorded HFOs (Fig 3C to E). None of the Ts65Dn controls showed HFOs (Fig 3B).

**Figure 3:**
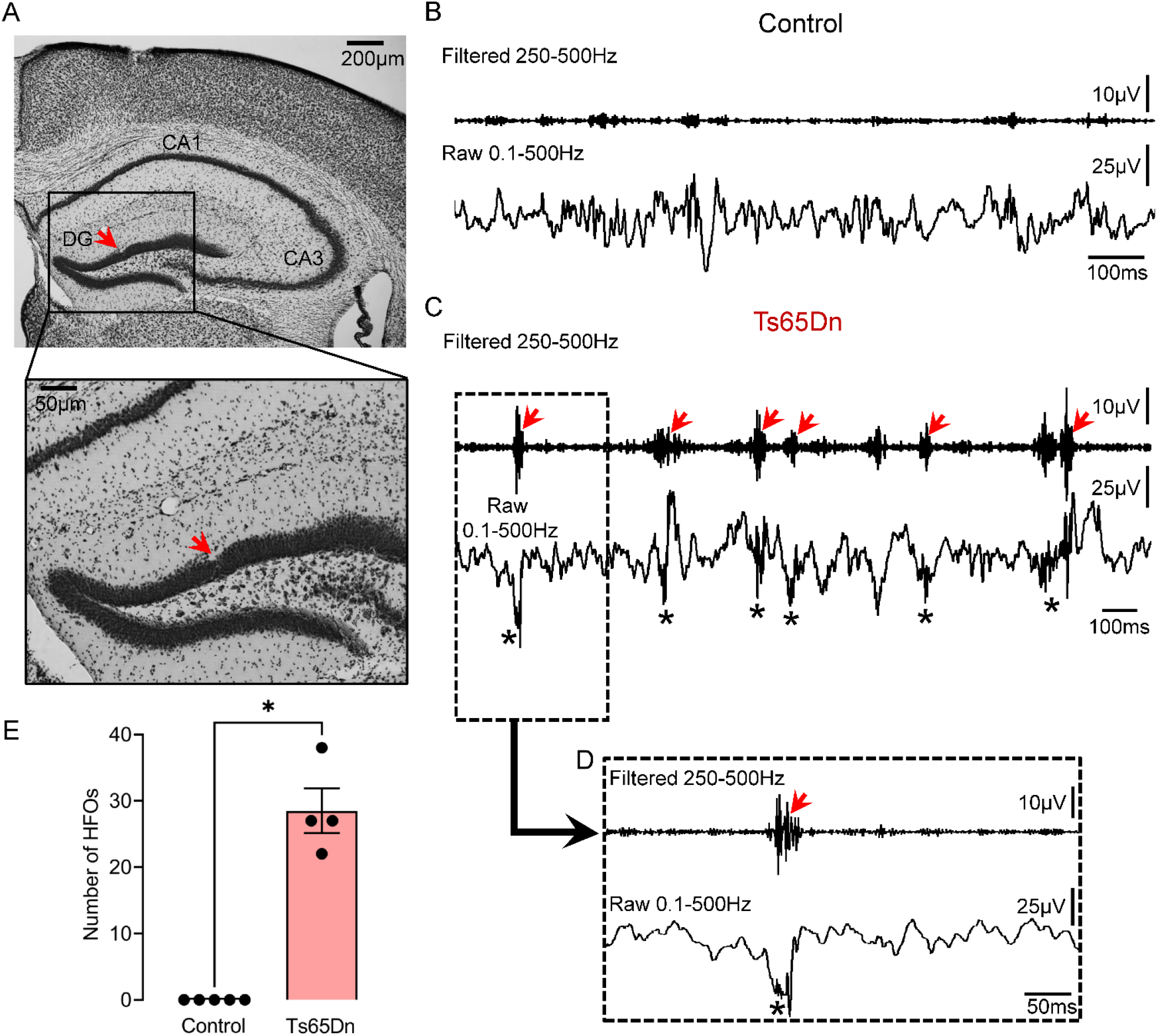
HFOs in the dentate gyrus of a mouse model of Down’s syndrome. (A) A representative Nissl-stained section confirming electrode placement in the DG (red arrow). Electrode targeting for both the control and transgenic mice is provided in Supplemental figure 4. (B) A control EEG recording from the DG during SWS is shown. No HFOs were detected. (C) Same as in B but the mouse was a Ts65Dn transgenic. Note HFOs in filtered (red arrows) and raw (asterisks) traces. (D) Expanded traces from the inset in C are shown. Note an HFO in the filtered trace (red arrow) which is also visible in the raw trace (asterisk). (E) Total number of HFOs recorded from control (n=5) and Ts65Dn transgenic (n=4) mice. Data from all behavioral states were pooled. The total number of HFOs was significantly greater in Ts65Dn vs. control mice (Mann-Whitney U-test, U=0, p=0.01).

### II. HFOs in AD mice are most frequent during slow wave sleep

We next analyzed HFOs separately for periods of wakefulness and sleep. Wakefulness included periods of awake immobility and exploration. Sleep was divided into SWS and REM based on theta/delta ratio as previously.^20^ Representative examples of HFOs and theta/delta ratio during sleep are shown in Fig 4A (Top panel).

**Figure 4:**
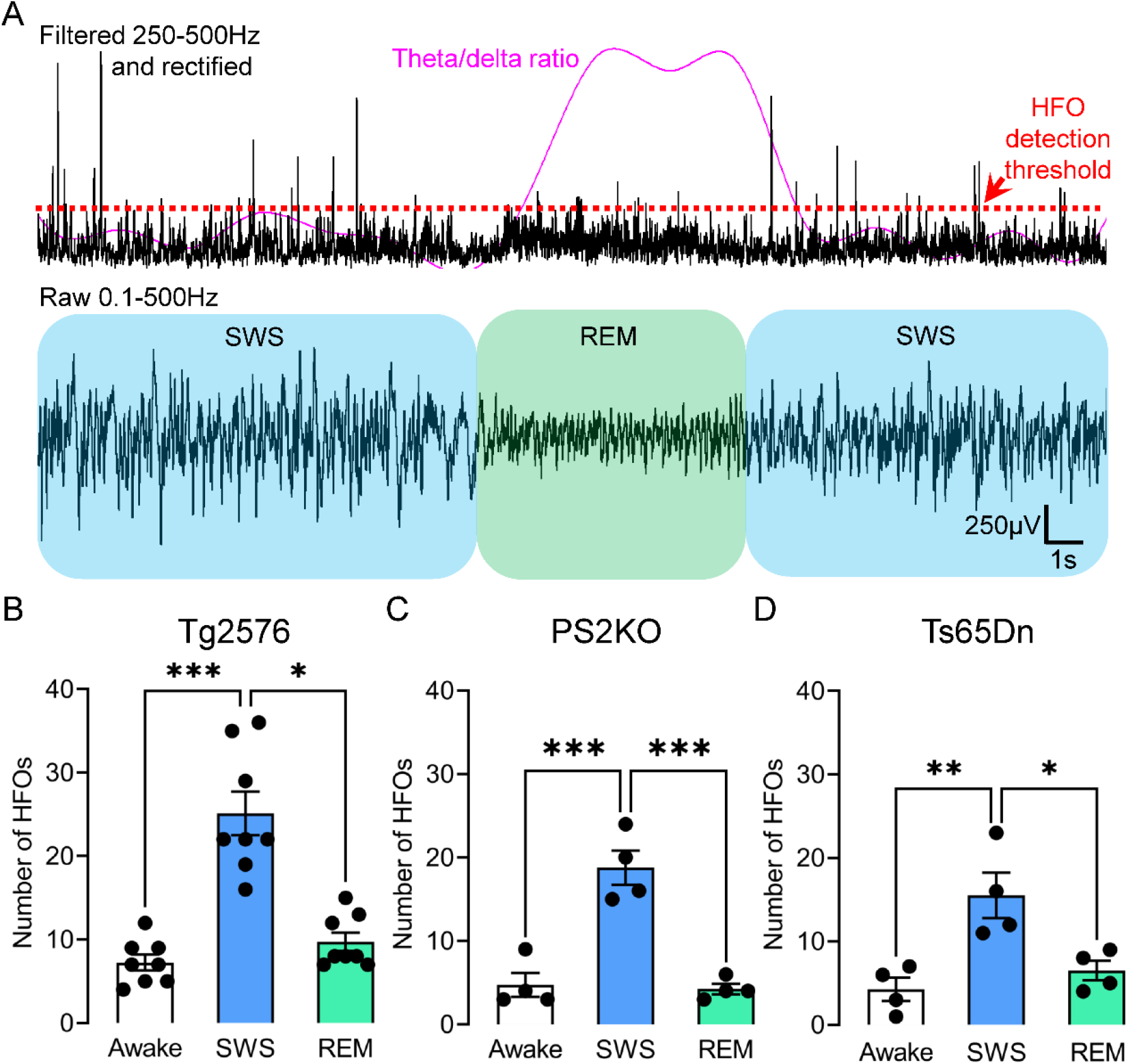
The number of HFOs in AD mouse models is elevated during slow wave sleep. (A) Top trace is filtered in the 250-500Hz frequency, rectified to show HFOs during a representative sleep period that includes both SWS (blue) and REM (green). Detection threshold for HFOs is denoted by a red dotted line and theta/delta ratio by a purple line. Bottom trace shows raw EEG during SWS (blue) and REM (green). Note frequent HFOs during SWS and reduced number of HFOs during REM, which corresponds to an increased theta/delta ratio. (B) Number of HFOs during wakefulness, SWS and REM sleep in Tg2576 transgenic mice (n=8). One-way ANOVA revealed a significant difference (Kruskal-Wallis test, H(3)=16.56, p=0.0003) and *post-hoc* tests showed a significantly greater number of HFOs during SWS vs. REM (p=0.01) or wakefulness (p=0.0003). There was no significant difference between the mean number of HFOs in wakefulness vs. REM (p=0.90). (C) Same as in B but for PS2KO transgenic mice (n=4). One-way ANOVA revealed a significant difference (F(2, 9)=30.39, p<0.0001). *Post-hoc* tests showed a significantly greater number of HFOs during SWS vs. REM (p=0.0002) or wakefulness (p=0.0002), but not between wakefulness and REM (p=0.81). (D) Same as in C but for Ts65Dn transgenic mice (n=4). One-way ANOVA revealed a significant difference (F(2, 9)=9.90, p<0.01). *Post-hoc* tests showed a significantly greater number of HFOs during SWS vs. REM (p=0.02) or wakefulness (p=0.006), but not wakefulness and REM (p=0.80).

Next, we compared HFOs between all 3 behavioral states. In Tg2576 mice (Fig 4B), one-way ANOVA showed that total HFO number was significantly different (Kruskal-Wallis test, H(3)=16.56, p=0.0003). *Post-hoc* tests showed a significantly greater number of HFOs during SWS vs. REM (p=0.01) or wakefulness (p=0.0003). Differences in the mean number of HFOs between wakefulness and REM were not significant (p=0.90).

In PS2KO mice (Fig 4C), behavioral states were also significantly different (F(2, 9)=30.39, p<0.0001). Like Tg2576 mice, *post-hoc* tests in PS2KO mice showed a significantly greater number of HFOs during SWS vs. REM (p=0.0002) or wakefulness (p=0.0002), but not wakefulness and REM (p=0.81).

Behavioral states were also different in Ts65Dn mice (F(2,9)=9.90, p=0.005; Fig 4D). Like the other mouse models, *post-hoc* tests showed a significantly greater number of HFOs during SWS vs. REM (p=0.02) or wakefulness (p=0.006), but not wakefulness and REM (p=0.80).

### III. Dentate gyrus HFOs are remarkably similar in spectral frequency and duration across different AD mouse models

In all AD models, time-frequency analysis revealed a well-localized HFO power (“frequency island”) in the 250-500Hz frequency range (Fig 5A to C). That frequency island was restricted to the time the HFO power peaked and thus coincided with an HFO in the filtered trace (Fig 5A to C; middle panels). To determine whether there are potential differences between models across behavioral states, a two-way ANOVA was used with model and behavioral state as factors (Fig 5D). Both model (F(2, 39)=8.45, p=0.0009) and behavioral state (F(2, 39)=49.86, p<0.0001) were significant; no significant interaction was found (F(4, 39)=1.21, p=0.32). *Post-hoc* comparisons showed that Tg2576 mice had a significantly greater number of HFOs vs. PS2KO (p=0.04) or Ts65Dn (p=0.001) mice. Although the total number of HFOs (all behavioral states combined, “All”; Fig 5D) was significantly different between models (One-way ANOVA, F(2, 13)=4.97, p=0.02), neither HFO duration (One-way ANOVA, F(2, 13)=0.54, p=0.59; Fig 5E) nor spectral frequency (Kruskal-Wallis test, H(3)=0.90, p=0.66; Fig 5F) was.

**Figure 5:**
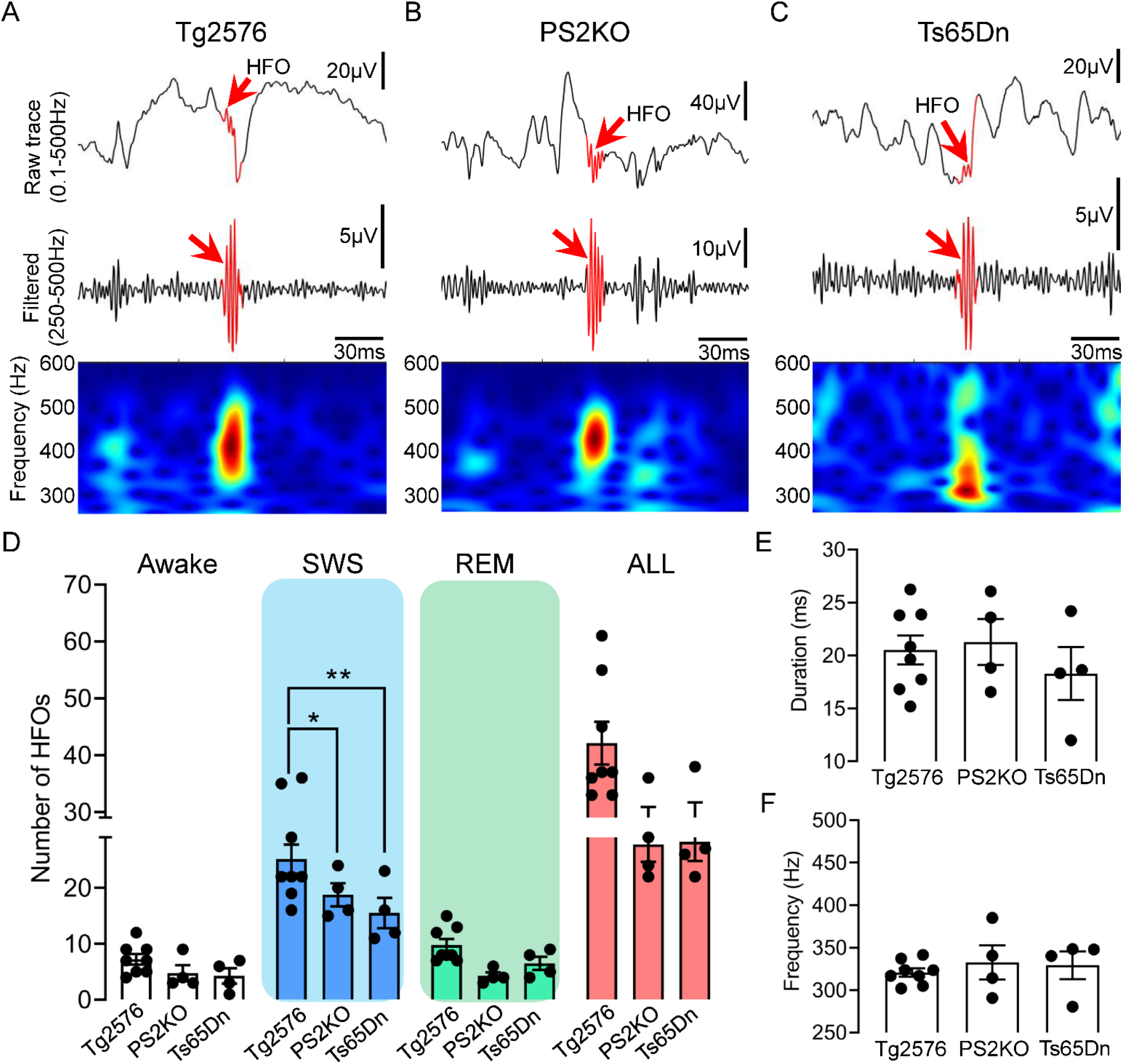
Comparison of spectral frequency, number and duration of HFOs between different AD mouse models. (A) Representative example of an HFO (red arrow) recorded in the DG of a Tg2576 transgenic mouse. Top trace is wideband recording (0.1-500Hz). The center is a filtered trace (250-500Hz), and at the bottom are spectral properties of the HFO in the 250-600Hz time-frequency domain. (B) Same as in A but for an HFO recorded from a PS2KO transgenic mouse. (C) Same as in B but for an HFO recorded from a Ts65Dn transgenic mouse. (D) Number of HFOs during wakefulness, SWS, REM and all states combined (“All”) between different models. A two-way ANOVA with model and behavioral state as factors showed a significant effect for model (F(2, 39)=8.45, p=0.0009) and behavioral state (F(2, 39)=49.86, p<0.0001) with no significant interaction (F(4, 39)=1.21, p=0.32). *Post-hoc* comparisons showed that HFOs during wakefulness or REM were not significantly different between models (all comparisons; p>0.05). However, SWS HFOs were significantly different between Tg2576 vs. PS2KO (p=0.04) and Ts65Dn (p=0.001) mice. The total number of HFOs in all behavioral states (“All”), pooled, was significantly different between models (F(2, 13)=4.97, p=0.02) but *post-hoc* tests did not confirm statistically significant differences in the number of HFOs in Tg2576 vs. PS2KO (p=0.06), and Ts65Dn mice (p=0.07). (E) HFO duration during SWS between different mouse models. No significant differences were found between models (One-way ANOVA, F(2, 13)=0.54, p=0.59). (F) Spectral frequency of HFOs during SWS in different mouse models. No significant differences were found (Kruskal-Wallis test, H(3)=0.90, p=0.66).

### IV. HFOs in the AD mouse models are indistinguishable from HFOs in two animal models of epilepsy

We next asked whether HFOs in AD models were different from HFOs recorded in epileptic animals. To that end, we used 2 well established animal models of epilepsy where HFOs are known to be robust: the IHKA model^15^ and the PILO model.^11^ Examples of HFOs from both models are shown in Fig 6A to B. Similar to AD mouse models, HFOs in epileptic animals showed a well-localized HFO power in the 250-500Hz frequency band which corresponded to an HFO in the filtered trace (Fig. 6A to B; middle panels).

We analyzed HFO duration (Fig 6C) during SWS across 3 different HFO groups: AD, IHKA and PILO. The AD group included HFOs from all AD mouse models since no difference in duration or spectral frequency was identified (Fig 5E to F). One-way ANOVA revealed no statistically significant differences in the mean duration of HFOs between AD, IHKA and PILO groups (F(2, 23)=1.84, p=0.18). When spectral frequency was compared, no significant differences were found (F(2, 23)=1.66, p=0.21; Fig 6D).

**Figure 6:**
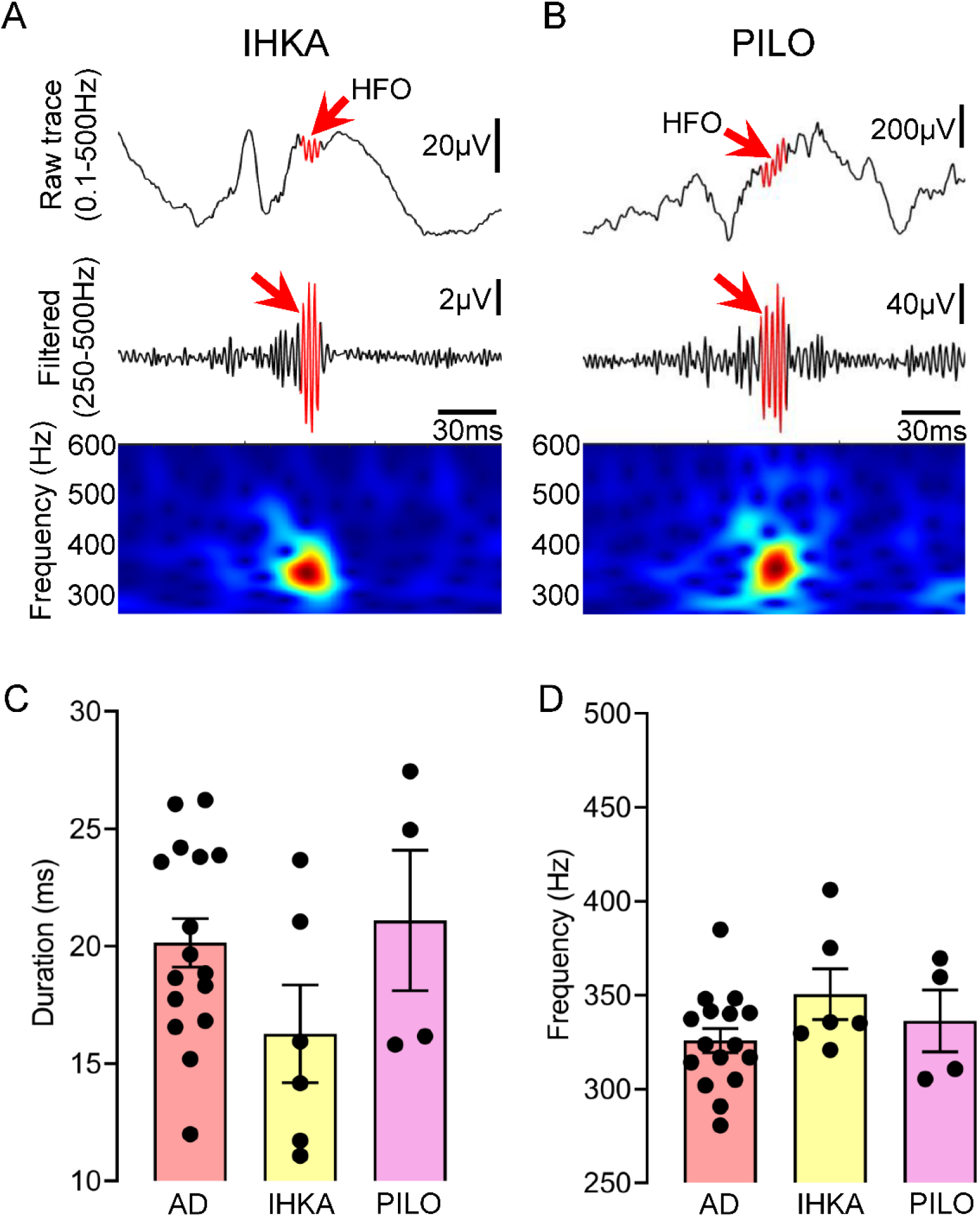
HFOs in the AD mouse models closely resemble HFOs in 2 animal models of epilepsy. (A) Representative example of an HFO (red arrow) recorded from the hippocampus of a mouse where kainic acid was injected. Top trace is wideband recording (0.1-500Hz). The center is filtered trace (250-500Hz), and bottom shows spectral properties of the HFO in the 250-600Hz time-frequency domain. (B) Same as in A but for an HFO recorded from the hippocampus of a PILO-treated mouse. (C) Comparison of HFO duration between AD, IHKA and PILO models. One-way ANOVA found no significant differences in HFO duration between models (F(2, 23)=1.84, p=0.18). (D) Same as in C but for spectral frequency. As in the case of HFO duration, one-way ANOVA found no significant differences between models (F(2, 23)=1.66, p=0.21).

### V. HFOs in interictal spikes (IIS) and detection of HFOs from a cortical contact

HFOs in epilepsy are known to be associated with IIS both *in vitro*,^33^ *in vivo*^34^ and in recordings from epilepsy patients.^35^ Thus, we asked if AD HFOs were also associated with IIS because IIS are common in patients with AD.^7^ To that end, we detected IIS (Fig 7A) and HFOs during REM sleep – a time when IIS are frequent in the Tg2576 mouse.^20^ Remarkably, the IIS we detected were associated with HFOs (Fig 7B). HFOs occurred before the peak amplitude of an IIS (Fig 7C), during an IIS (Fig 7D) or following an IIS’ peak (Fig 7E). IIS HFOs showed a well-localized frequency island in the time-frequency domain (Fig 7C to E; bottom panels) resembling frequency islands of isolated HFOs (Fig 5A to C; bottom panels). We found that HFOs were significantly shorter in duration when associated with IIS vs. isolated HFOs (paired Student’s t-test, t=3.33, df=4, p=0.02; Fig 7F). Also, IIS HFOs showed significantly higher spectral frequency compared to HFOs occurring independent of IIS (paired Student’s t-test, t=3.01, df=4, p=0.03; Fig 7G).

**Figure 7:**
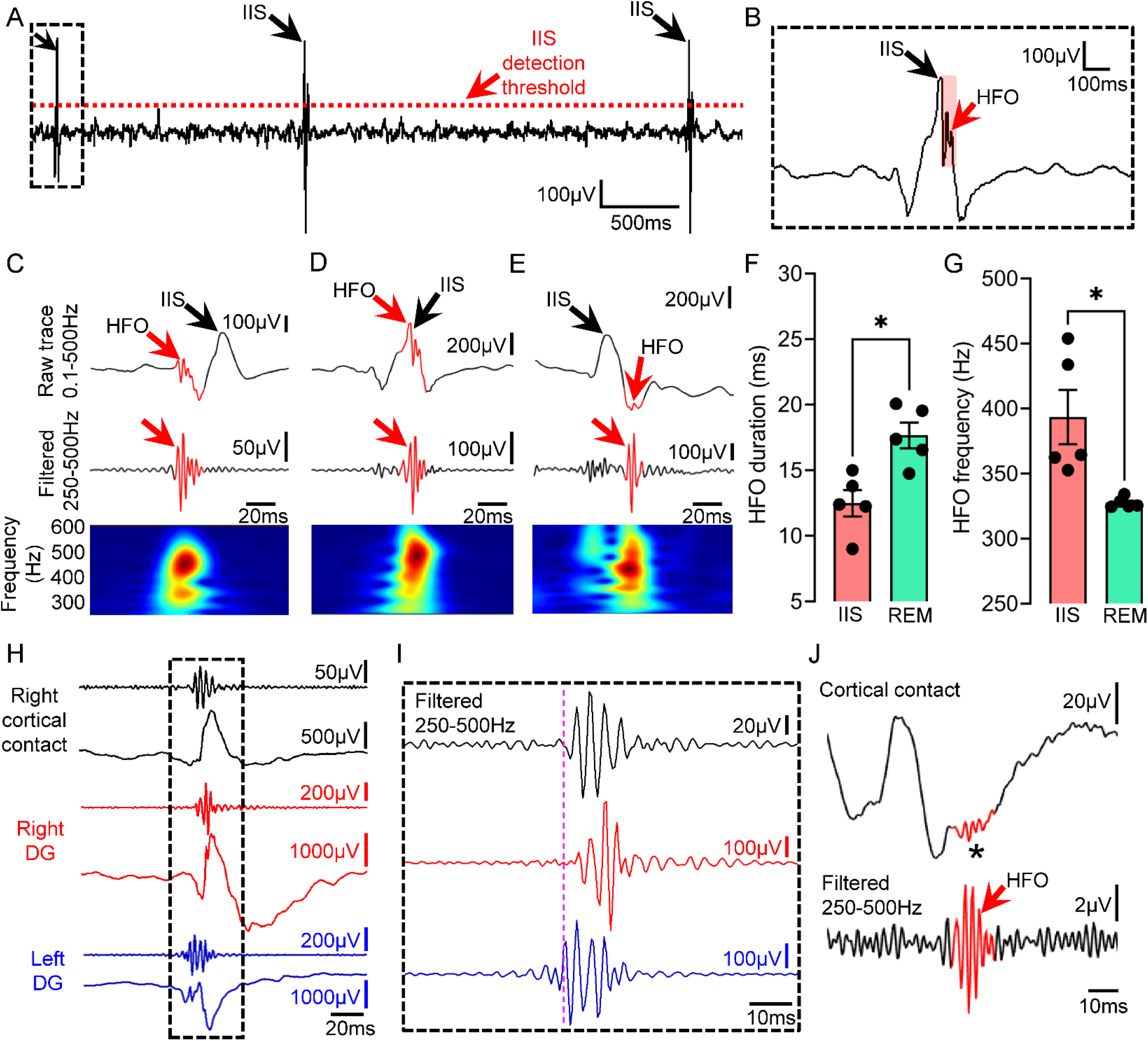
HFOs associated with IIS and detection of HFOs from the cortical surface. (A) Detection of IIS during REM sleep in a Tg2576 transgenic mouse. Threshold used for IIS detection is denoted by a red dotted line. (B) Expanded trace of a representative IIS from the inset in A. Note the presence of an HFO (red arrow). (C) Example of an HFO occurring before the peak amplitude of an IIS. (D) Example of an HFO occurring during an IIS. (E) Example of an HFO occurring after an IIS peak. (F) HFOs associated with IIS were significantly shorter in duration compared to HFOs occurring independent of IIS (paired Student’s t-test, t=3.33, df=4, p=0.02). (G) Same as in F but for spectral frequency. HFOs associated with IIS showed significantly higher spectral frequency vs. HFOs independent of IIS (paired Student’s t-test, t=3.01, df=4, p=0.03). (H) HFOs during an IIS recorded simultaneously from left (blue traces) and right DG (red traces) and overlying right frontal cortex (black traces) using a subdural screw electrode. (I) Expanded traces from the inset in H showing filtered traces in the 250-500Hz frequency. Note that HFOs do not start or peak at the same time in all recording channels. Vertical purple line denotes onset of the HFO in the left DG. (J) Representative example of an HFO (red arrow) recorded with a subdural screw electrode from the right frontal cortex during SWS. Note presence of the HFO in the unfiltered trace (asterisk).

Notably, simultaneous intra-hippocampal and cortical recordings showed that HFOs can be detected from the cortical surface. Fig 7H shows an IIS recorded from left and right DG, and right cortical contact. Note the presence of HFOs in all recording sites and asynchronous time of onset between sites (Fig 7I; purple dotted line). We also detected cortical HFOs occurring in isolation (independent of IIS) during SWS and a representative example is shown in Fig 7J.

### VI. HFOs in the AD mouse models occur during seizures

As seizures are common in familial AD^4^ and animal models of epilepsy^11,15^ we asked whether seizures in AD are accompanied by HFOs as they are in animal models of epilepsy^11,15^ and epilepsy patients.^36^ To that end, we analyzed HFOs during a representative seizure recorded from the DG of a 18-month-old Tg2576 transgenic mouse (Fig 8A). The seizure was recorded simultaneously from left and right DG, and it was a Racine stage 5 convulsive seizure.^26^ HFOs were detected in both left and right DG and were characterized by spectral frequency exceeding 200Hz (Fig 8B). We found that HFOs during seizures were associated with different spike morphologies (Fig 8B1 vs. 8B2) and were also detected between spikes (Fig 8B3; left DG).

**Figure 8:**
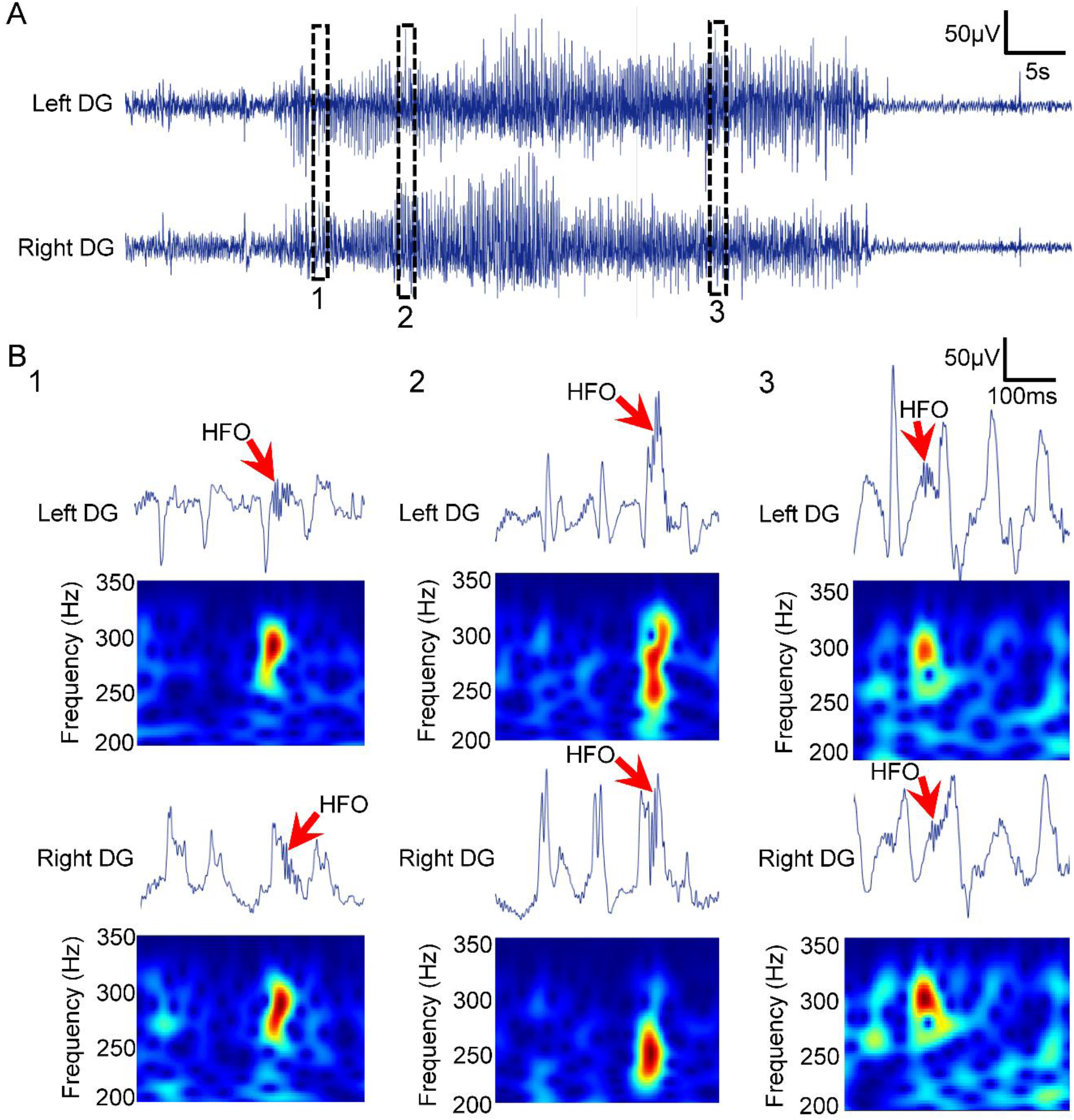
Spectral properties of HFOs during a representative seizure in a Tg2576 mouse. (A) Example of a convulsive seizure recorded from the DG of an 18-month-old Tg2576 transgenic mouse. Top trace is recording from the left DG and bottom is from the right DG. Insets 1, 2 and 3 are expanded in B. (B) Spectral frequency of HFOs during a convulsive seizure. Top trace is wideband recording. Bottom panel shows spectral properties of the HFOs with frequency >200Hz in the time-frequency domain. B1: An HFO occurring just after an IIS in left and right DG. B2: An HFO during an IIS in the left and the right DG. B3: An HFO at the time between 2 IIS in the left DG and during an IIS in the right DG.

## DISCUSSION

This study was undertaken to determine whether HFOs can be detected in AD mouse models. Using 3 mouse models of AD, each with a different mutation and neuropathology, we have demonstrated that HFOs between 250 and 500Hz are a novel electrophysiological disturbance. Notably, we found that most HFOs occur during SWS but durations and the frequency islands’ were not significantly different between behavioral states. Also, HFOs could be recorded by intra-hippocampal electrodes as well as the cortical surface, making non-invasive recordings feasible. There also were striking commonalities with HFOs in animal models of epilepsy. Last, HFOs could occur in conjunction with other epileptiform activity such as IIS and seizures, or in isolation, and HFO characteristics depended on this timing.

### HFOs as a novel EEG abnormality in AD

We defined HFOs as oscillations >250Hz in line with previous studies^14,15^ and consider them abnormal, consistent with human epilepsy studies of HFOs >250Hz showing they were absent from normal hippocampus and therefore pathological.^37^ We recorded robust HFOs from the DG, where there normally are faster oscillations relative to other areas. However, the fast activity is characterized by frequencies <150 Hz.^38^ Therefore it is easily distinguished from HFOs >250Hz. Recordings from overlying area CA1 did not show HFOs at the time of DG HFOs. However, we cannot exclude the possibility that other hippocampal areas such as CA3 contribute to DG HFOs. Taken together with the lack of HFOs in age-matched controls of the 3 different AD mouse lines; our data suggest that HFOs in the DG of AD mice represent an abnormal electrical pattern.

We recorded HFOs as early as 1month in the DG of Tg2576 transgenic mice. The DG is known to exhibit amyloid-β plaque long after 1month,^39^ but its excitability is increased by approximately 1month as shown by IIS.^20^ Robust occurrence of HFOs as early as 1month indicates that HFOs and IIS are likely to be the first electrophysiological disturbances occurring in the AD mouse model. Note that seizures in Tg2576 mice occur by 2months, but not at 1month of age.^20^ Therefore, HFOs and IIS occur before seizures, which is interesting because in epilepsy a spontaneous seizure often has HFOs and IIS before it.^11,15,34^ Taken together, the presence of HFOs when there is no epilepsy or robust AD neuropathology suggests that HFOs are not intrinsic to either condition, but they rather reflect an atypical EEG pattern that may accompany both.

### Brain state dependency of HFOs in AD mouse models

We found most HFOs occur in SWS, consistent with the observations that EEG abnormalities in sleep occur during SWS in humans.^18^ A greater number of HFOs during SWS is consistent with human^28^ and rodent^27^ epilepsy data showing that HFOs in epilepsy occur primarily during SWS. HFOs in humans are known to be facilitated by the presence of slow waves during SWS^40^ similar to the slow wave activity we recorded HFOs on in our mice. One of the reasons why HFOs are facilitated during SWS could be related to periods of high synchronization such as those during transitions from depolarizing ‘up’ states to hyperpolarzing ‘down’ states.^40^

### Similarities to epileptic animals

Currently, there is no evidence that HFOs similar to those recorded in epilepsy can be recorded in the context of other neuropsychiatric disorders. Therefore, finding HFOs in AD mouse models and finding they are similar to the HFOs in epilepsy models was significant. Both spectral frequency and duration of HFOs were remarkably similar, suggesting that HFOs possibly reflect the same electrical abnormality with similar underlying mechanisms. Although still debated, one mechanism is that HFOs are due to synchronous firing of adjacent principal (e.g., granule) neurons at slightly different times.^41^ The times occur without much of an interval, so the extracellular manifestation is a rapid oscillation (HFO). The timing also is so fast that synaptic mechanisms are unlikely. Instead, gap junctions,^42^ intrinsic properties and ion channels^43^ have been suggested to contribute, and there are other possibilities. In AD models, altered intrinsic properties^21^ and ion channels^16,44^ have been identified already which may contribute to HFOs.

### HFOs in IIS and seizures in AD

HFOs occurred during IIS in sleep. Their spectral frequency was increased, and duration was decreased compared to isolated HFOs. Such differences in HFOs are reminiscent of HFOs during IIS in seizure-generating areas with epilepsy pathology compared to remote areas to the seizure-generating zone.^45^ Thus it is possible that co-occurrence of HFOs with IIS in AD mice reflect underlying pathophysiological processes responsible for increased excitability and/or neuropathology. Indeed, clusters of hyperactive neurons are known to surround amyloid-β plaques in experimental AD.^46^ Thus, HFOs with IIS could help identifying brain areas with increased excitability and neuropathology in AD. Nevertheless, future studies looking at co-occurrence of IIS and HFOs beyond REM sleep are important because we found that HFOs are also detectable during other behavioral states besides REM sleep.

HFOs were also detected during seizures. HFOs during seizures in epilepsy are particularly useful for mapping brain areas where seizures initiate.^12^ Likewise, HFOs during seizures in AD may also be helpful in defining seizure-prone areas. Regardless, the association of HFOs with seizures in AD models further supports the idea that the HFOs we recorded reflect abnormal electrical patterns. However, HFOs during IIS, seizures or isolated HFOs might not share similar characteristics. Thus, they may differ in spectral frequency or duration. HFOs with a faster spectral frequency may be due to changes in intrinsic properties, such as action potentials with shorter half-width, steeper rate of rise, or faster rate of decay. Seizures and IIS may also exert effects on HFOs, influencing the characteristics of the HFOs.

### Potential role of HFOs in monitoring disease progression in AD

HFOs in epilepsy are best known as “spatial” biomarkers of epileptogenic tissue.^12,14^ However, beyond their important localizing value there is some data suggesting that the number of HFOs at a certain time during the disease reflects disease severity.^47^ Indeed, non-invasive recordings of HFOs from the scalp suggested that scalp-recorded HFOs are greater for larger areas of damaged tissue.^48^ Notably, we did not find a statistically significant correlation between the total number of HFOs and age of transgenic mice (Pearson r=-0.23, p=0.37, n=16; Supplemental figure 1A). Instead, we found a non-linear relationship between the total number of HFOs and age of transgenic mice (Supplemental figure 1B).

In the present study, we found more HFOs in Tg2576 mice which have the most plaque pathology.^19^ Notably, differences in HFO occurrence between models were robust in SWS but not during wakefulness or REM, which is interesting because SWS is the sleep stage when amyloid-β is being cleared from the brain.^49^ If HFOs disrupt SWS in AD, then it is possible that their robust occurrence during SWS contributes to increased accumulation of amyloid-β over time. These findings also suggest that SWS could be a preferred state of monitoring when AD HFOs are investigated, resulting in a more efficient evaluation of people at risk for AD.

### Could HFOs help predict epilepsy development, AD development or both?

Tg2576 mice that develop seizures after 7weeks of age had HFOs at 1month of age. HFOs also occurred in PS2KO and Ts65Dn mice where seizures haven’t been reported. Thus, it is not necessary that animals develop epilepsy in AD models to show HFOs, at least in the mouse models we examined. It also is not necessary that animals show robust signs of AD neuropathology, i.e., amyloid-β plaques, to show HFOs because Tg2576 mice had HFOs prior to the age when amyloid-β deposits are first detected, 6months.^19^ Furthermore, PS2KO and Ts65Dn mice had HFOs but do not develop amyloid-β plaques.^24^ Taken together, the presence of HFOs when there is no epilepsy or robust AD neuropathology suggest that HFOs are an EEG abnormality that is not necessarily linked to either AD or epilepsy, at least in animal models.

## CONCLUSIONS

Revealing DG and cortical HFOs in AD mouse models suggest a new EEG abnormality in AD with important translational value. The occurrence of HFOs during IIS and seizures suggests that HFOs are related to hyperexcitability in AD, supporting accumulating evidence that increased excitability is a contributor to the disease. However, at a late stage, the degeneration caused by a history of overactivity is likely to depress synaptic function, as the majority of those who study AD have found (discussed elsewhere).^20^ The association of HFOs with sleep provides a new reason for sleep disturbance in AD which could contribute to disrupted sleep functions such as impaired memory consolidation and lower amyloid-β clearance. The mechanisms proposed to underlie HFOs in epilepsy, and the common characteristics of HFOs in AD mouse models and epilepsy models, suggest changes in pyramidal cell firing are an important part of AD pathophysiology, and indeed ion channels^16,44^ underlying firing have been suggested but amyloid-β and hyperphosphorylated tau remain the most common focus for treatment.

This study also suggests that HFOs could be a new biomarker in AD. The ability to record HFOs and epileptiform activity from different brain areas, and do so non-invasively,^50-52^ strengthens that view. Additional benefits include the ability to conduct EEG recordings repeatedly over time to characterize disease progression. The safety of EEG also allows recording at younger ages than current biomarkers allow, which could provide new insight into the earliest phases of AD.

## Abbreviations

AD: Alzheimer’s disease
HFOs: High frequency oscillations
IHKA: Intra-hippocampal kainic acid
IIS: Interictal spike
PILO: Pilocarpine
PS2KO: Presenilin 2 knock-out
REM: Rapid eye movement sleep
SWS: Slow wave sleep

## ACKNOWLEDGEMENTS

We would like to thank Dr. Ralph Nixon for providing the PS2KO mice and Dr. Stephen Ginsberg for providing the Ts65Dn mice. We thank Drs. Azahara Oliva and Sam Mckenzie for initial methodological help with silicon probe recordings. We also thank John LaFrancois for technical assistance with the PILO model. This project was supported by NIH grants R01 AG-055328 and R01 NS-106983, and the New York State Office of Mental Health.

## AUTHOR CONTRIBUTIONS

Conceptualization: CPL, HES

Data acquisition: CPL

Methodology: CPL

Data analysis: CPL, HES

Funding acquisition: HES

Resources: HES

Project administration: HES

Supervision: HES

Data visualization: CPL, HES

Writing – original draft: CPL

Writing – review & editing: CPL, HES

## CONFLICTS OF INTEREST/ETHICAL PUBLICATION STATEMENT

None of the authors has any con?ict of interest to disclose. We con?rm that we have read the Journal’s position on issues involved in ethical publication and affirm that this report is consistent with those guidelines.

## DATA AVAILABILITY STATEMENT

Data will be available from the corresponding author upon request.

## Supporting Information

**Supplemental table 1 legend:**
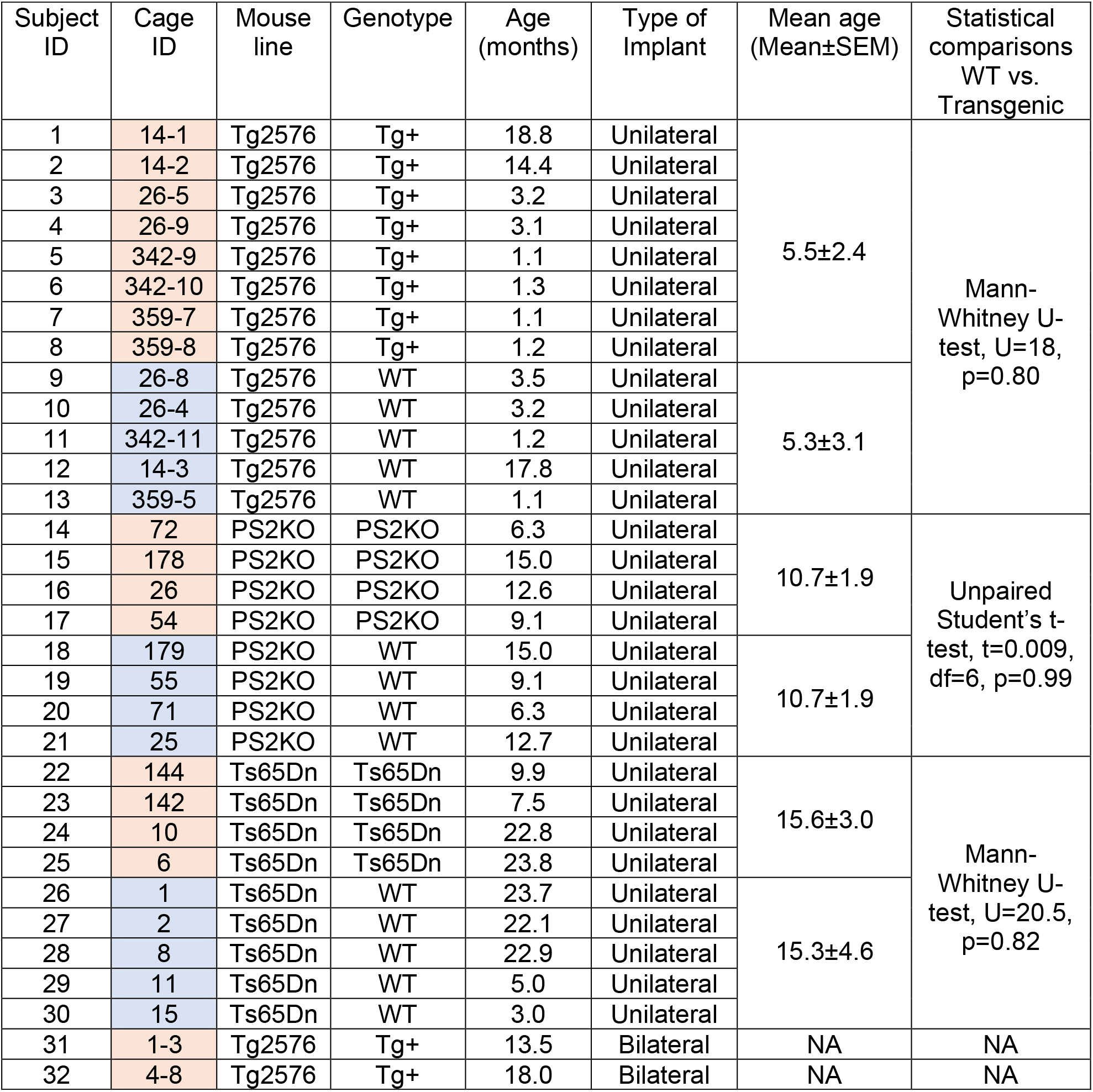
Animals used in the study. For each animal we note the subject ID, cage ID, mouse line (Tg2576, PS2KO, Ts65Dn), genotype, age (months), type of implant (unilateral, bilateral), mean age and statistical comparisons between the age of wild type (WT) and transgenic mice. Orange shading denotes transgenic mice and blue WT mice. <u>Tg2576 mouse line:</u> For the Tg2576 mouse line we used n=8 transgenic and n=5 WT mice. The mean age of transgenic mice was 5.5±2.4months and for the WT mice 5.3±3.1months. There was no significant age difference between transgenic and WT mice (Mann-Whitney U-test, U=18, p=0.80). Subjects #31 and #32 were used for seizure monitoring. <u>PS2KO mouse line:</u> For the PS2KO mouse line we used n=4 transgenic and n=4 WT mice. The mean age of transgenic mice was 10.7±1.9months and for the WT mice 10.7±1.9months. There was no significant age difference between transgenic and WT mice (unpaired Student’s t-test, t=0.009, df=6, p=0.99). <u>Ts65Dn mouse line:</u> For the Ts65Dn mouse line we used n=4 transgenic and n=4 WT mice. The mean age of transgenic mice was 15.6±3.0months and for the WT mice 15.3±4.6months. There was no significant age difference between transgenic and WT mice (Mann-Whitney U-test, U=20.5, p=0.82). Also, there were no statistically significant differences in age between the controls of different mouse lines (Kruskal-Wallis test, H(3)=3.89, p=0.14) and no differences in transgenics of the different mouse lines (Kruskal-Wallis test, H(3)=4.84, p=0.08).

Lack of correlation between the number of HFOs and age

**Supplemental figure 1:**
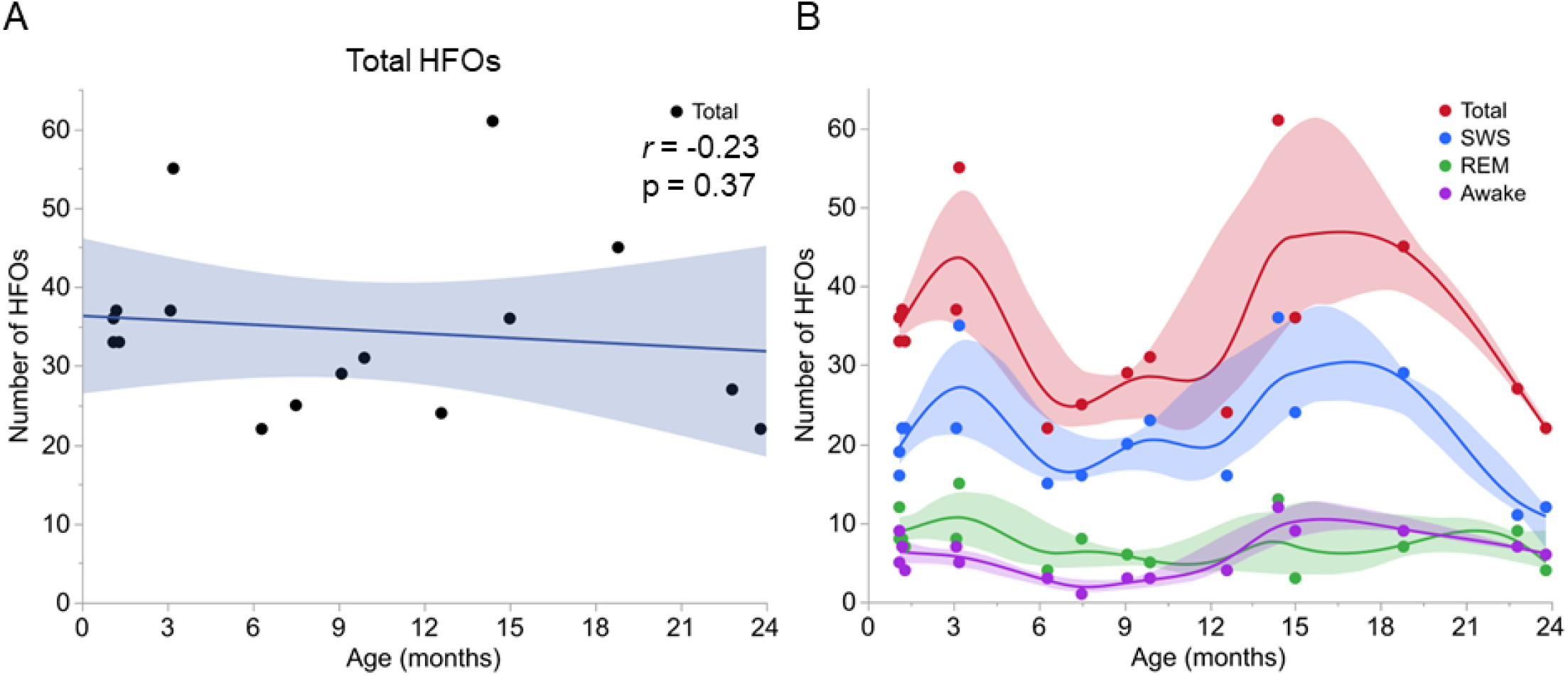
Lack of correlation between the total number of HFOs and age. (A) The total number of HFOs and age of transgenic mice is shown (Tg2576, n=8; PS2KO, n=4; Ts65Dn, n=4). The shading reflects the 95% confidence interval. We found no statistically significant correlation between the total number of HFOs and age of transgenic mice (Pearson r=-0.23, p=0.37, n=16). (B) Same as in A, but for HFOs recorded during SWS, REM, wakefulness, or total number of HFOs in all behavioral states combined. Note the non-linear relationship between the number of HFOs and age of transgenic mice (Tg2576, n=8; PS2KO, n=4; Ts65Dn, n=4). The shading reflects the 95% confidence interval.

Histological verification of electrode placement

**Tg2576 mouse line**

**Supplemental figure 2:**
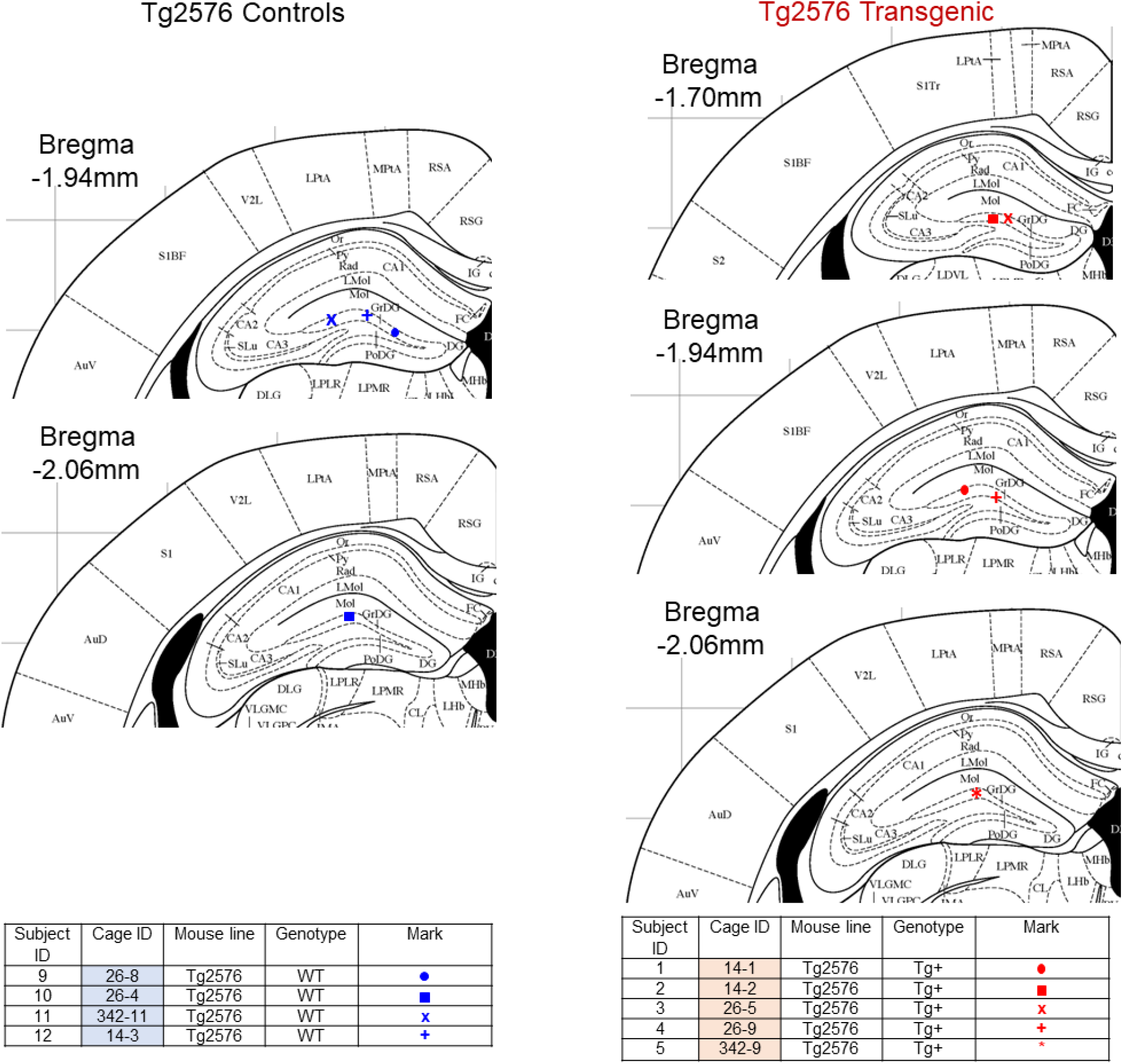
Verification of electrode placement in Tg2576 mice. Plots show placement of electrodes in each mouse used in the study. Control mice (n=4) are shown on the left and transgenic mice (n=5) on the right. The list of animals that electrodes placement was verified is provided below each plot. Different marks are used for each animal. Blue marks denote controls and red marks transgenic mice. The coronal diagrams are from a common mouse stereotaxic atlas.^1^

**PS2KO mouse line**

**Supplemental figure 3:**
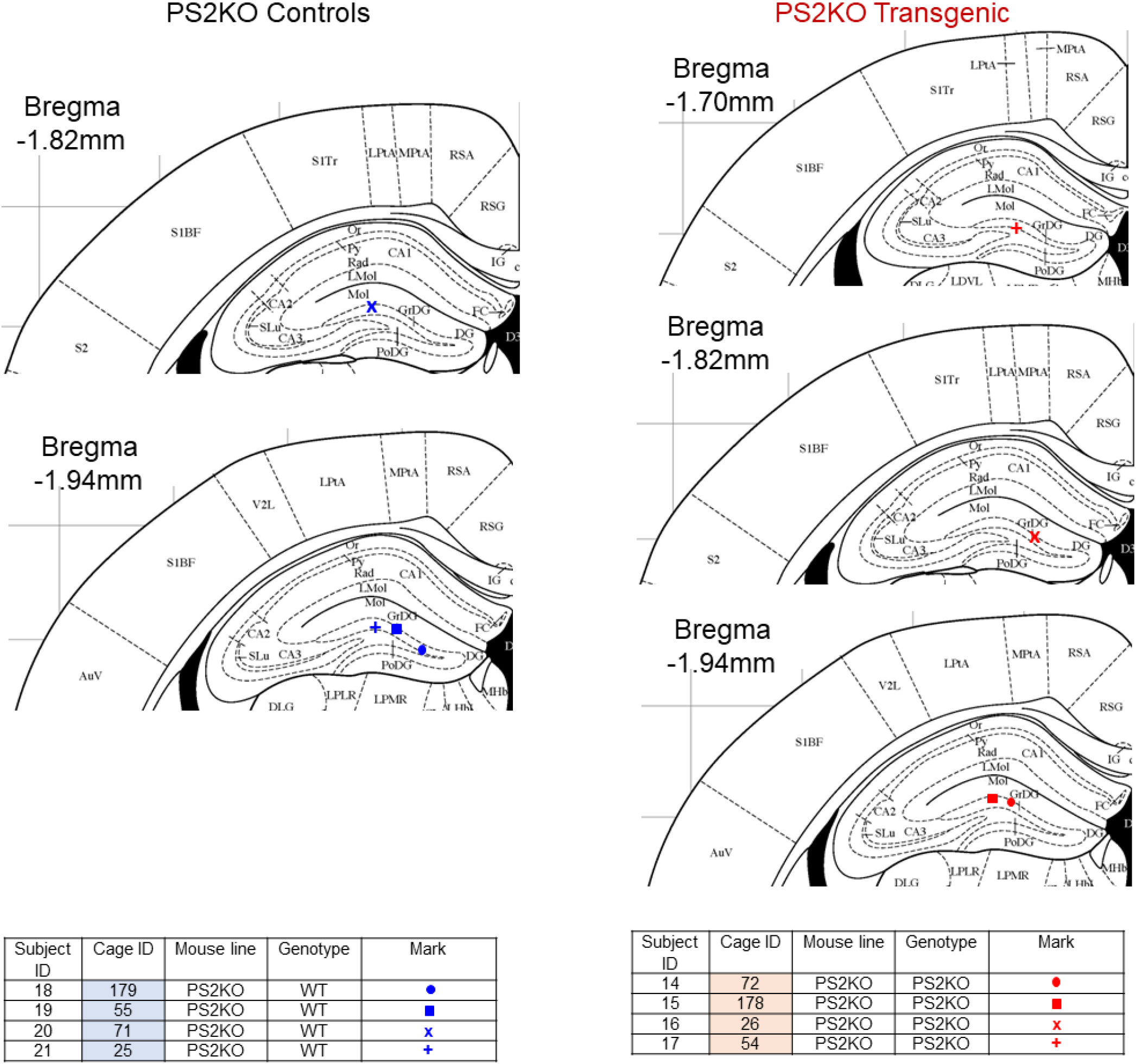
Verification of electrode placement in PS2KO mice. Plots show placement of electrodes in each mouse used in the study. Control mice (n=4) are shown on the left and transgenic mice (n=4) on the right. The list of animals that electrodes placement was verified is provided below each plot. Different marks are used for each animal. Blue marks denote controls and red marks transgenic mice. The coronal diagrams are from a common mouse stereotaxic atlas.^1^

**Ts65Dn mouse line**

**Supplemental figure 4:**
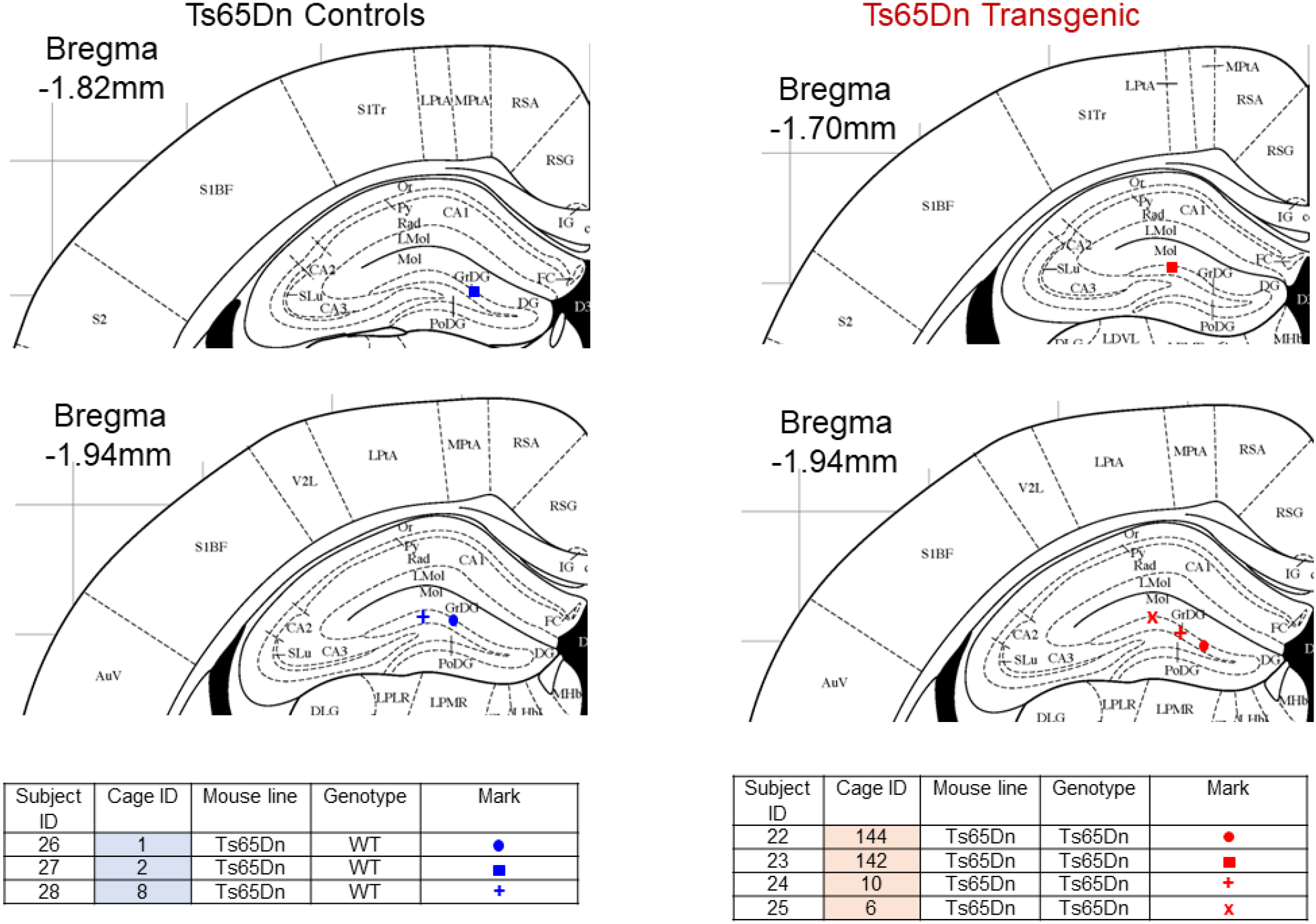
Verification of electrode placement in Ts65Dn mice. Plots show placement of electrodes in each mouse used in the study. Control mice (n=3) are shown on the left and transgenic mice (n=4) on the right. The list of animals that electrodes placement was verified is provided below each plot. Different marks are used for each animal. Blue marks denote controls and red marks transgenic mice. The coronal diagrams are from a common mouse stereotaxic atlas.^1^

HFO occurrence across different recording days

**Supplemental figure 5:**
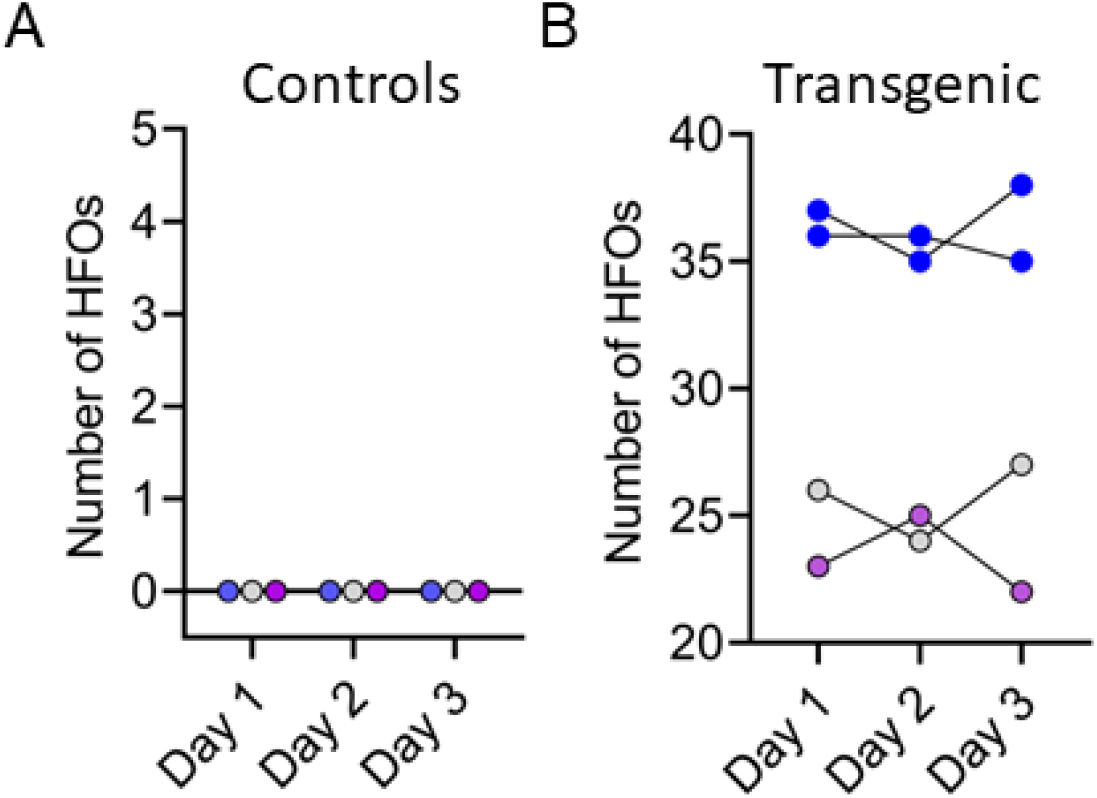
Number of HFOs during SWS in consecutive recording days. (A) Number of HFOs recorded in control mice (n=3) showed absence of HFOs in every recording day (Day 1, Day 2, Day 3). For this analysis, we used one mouse from each mouse line and the number of HFOs during SWS was analyzed during 10min periods of 3 consecutive days. Blue = Tg2576; Grey = PS2KO; Purple = Ts65Dn. (B) Number of HFOs recorded in transgenic mice (n=4) during 10min periods of SWS for 3 different recording days. No significant differences in the number of HFOs during SWS were found between recording days (one-way Repeated Measures ANOVA, F(2, 6)=0.1 5, p=0.87). Blue = Tg2576; Grey = PS2KO; Purple = Ts65Dn.

Layer-resolved profile of an HFO in AD

**Supplemental figure 6:**
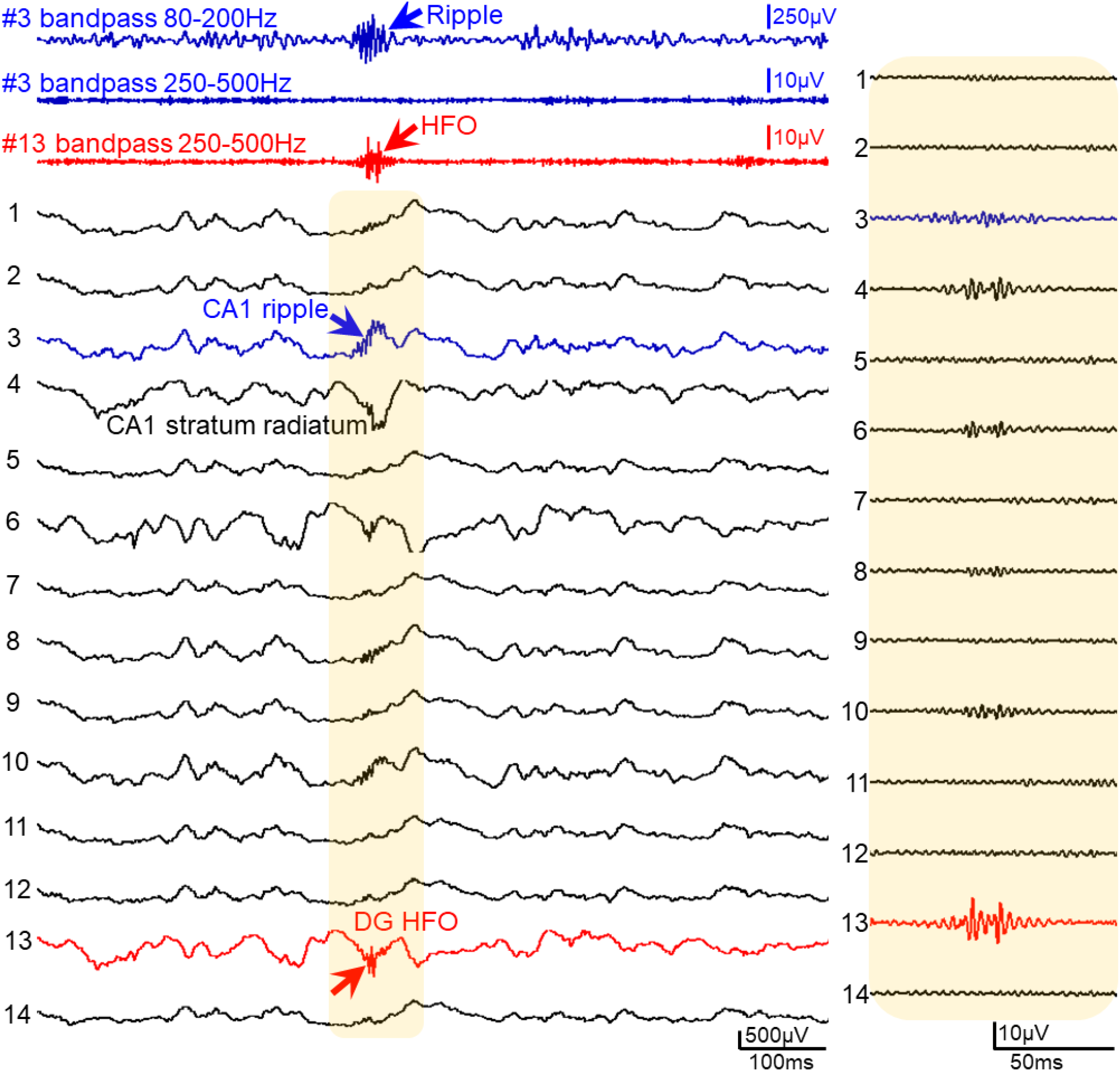
Layer-resolved profile of an HFO in AD. Representative layer-resolved profile of an HFO recorded in the DG in a Tg2576 mouse. A 16-channel silicon probe was used for the recording (2 channels were excluded from the analysis because of electrical noise). The spacing between electrodes is 100μm. Channels are organized from the most superficial one (channel #1) to the deepest (channel #14). Electrophysiological landmarks were used to determine the position of each electrode relative to cell layers. For example, the CA1 pyramidal layer was identified by gamma oscillations and ripples in the 80-200Hz frequency range.^2^ Channels in the DG were identified by the presence of fast activity in the gamma (∼80-150Hz) frequency range and in those channels the granule cell layer was identified by the presence of multiunit activity (not shown here). A DG HFO (red arrow) was recorded in channel #13 and a bandpass filtered trace of that channel is shown above in red (#13 bandpass 250-500Hz). Note robust HFO activity in the 250-500Hz frequency in the bandpass filtered channel #13 (red arrow). We also filtered the channel where ripple activity was robust (channel #3). That channel corresponded to the CA1 pyramidal layer. We found that there was no HFO activity in the 250-500Hz frequency range (#3 bandpass 250-500Hz). Instead, ripple activity (blue arrow) was robust when channel #3 was filtered in the 80-200Hz frequency range (#3 bandpass 80-200Hz). These results were similar for 7 silicon probe recordings from 5 transgenic mice (n= 2 Tg2576, n= 2 PS2KO, n= 1 Ts65Dn). The shaded area in yellow is expanded in time on the right to show filtered channels for HFOs in the 250-500Hz range. Note robust HFOs in the DG (red trace) and absence of HFOs from the CA1 and other channels.

## Notes

### Competing Interest Statement

The authors have declared no competing interest.

### Summary of Updates

1) New data were added in Figure 1. The new data included transgenic and control mice that were recorded at about 1month of age 2) The ages of all mice included in the study are now included in Supplemental table 1 along with other information 3) We examined possible correlation between the age of the mice and number of HFOs (Supplemental figure 1) 4) We provide electrode placement for mice used for histology (Supplemental figure 2-4) 5) We provide new data on HFO occurrence across different recording days (Supplemental figure 5) 6) We provide an example of a layer-resolved profile of an HFO in AD (Supplemental figure 5) 7) We added new text in the Discussion to discuss whether HFOs could help predict epilepsy development, AD development or both

